# Comparative single-cell trajectory network enrichment identifies pseudo-temporal systems biology patterns in hematopoiesis and CD8 T-cell development

**DOI:** 10.1101/2020.04.02.021295

**Authors:** Alexander G. B. Grønning, Mhaned Oubounyt, Kristiyan Kanev, Jesper Lund, Tim Kacprowski, Dietmar Zehn, Richard Röttger, Jan Baumbach

## Abstract

Single cell transcriptomics (scRNA-seq) technologies allow for investigating cellular processes on an unprecedented resolution. While software packages for scRNA-seq raw data analysis exist, no method for the extraction of systems biology signatures that drive different pseudo-time trajectories exists. Hence, pseudo-temporal molecular sub-network expression profiles remain undetermined, thus, hampering our understanding of the molecular control of cellular development on a single cell resolution. We have developed Scellnetor, the first network-constraint time-series clustering algorithm implemented as interactive webtool to identify modules of genes connected in a molecular interaction network that show differentiating temporal expression patterns. Scellnetor allows selecting two differentiation courses or two developmental trajectories for comparison on a systems biology level. Scellnetor identifies mechanisms driving hematopoiesis in mouse and mechanistically interpretable subnetworks driving dysfunctional CD8 T-cell development in chronic infections. Scellnetor is the first method to allow for single cell trajectory network enrichment for systems level hypotheses generation, thus lifting scRNA-seq data analysis to a systems biology level. It is available as an interactive online tool at https://exbio.wzw.tum.de/scellnetor/.

## Introduction

Single-cell RNA sequencing (scRNA-seq) allows researchers to perform cellular developmental studies with a hitherto unseen fine granularity. Single-cell transcriptomes have paved the way for novel discoveries in various biomedical fields by improving the understanding of how transcriptional profiles relate to cell phenotypes. A range of algorithms have been invented for clustering of scRNA-seq data and for inferring differentiation trajectories^1–3^. Clustering assumes that single cells can be divided into distinct groups whereas trajectory inference aims to arrange cells such that continuous phenotypes can be traced on a low dimensional cell map^4^. Important examples of the latter include diffusion maps^5,6^ and pseudotemporal ordering of single cells^3,7^. Both algorithms seek to position single cells such that their coordinates reflect their developmental statuses in relation to the other cells. Additionally, several software packages have been developed for the entire analysis pipeline, from pre-processing to clustering and identification of differentially expressed genes. Scanpy^8^, Seurat^9^ and SINCERA^10^ are examples of such software packages. Though scRNA-seq data is still challenged by noise^11^, combinations of different tools and algorithms have helped to unravel hidden inter-cellular mechanisms and shed light on unknown cellular paths of differentiation and disease progression^12–14^.

Typical computational analyses of single cell gene expression data involve a pre-processing step, where, e.g., cells with high levels of mitochondrial DNA and few expressed genes are removed^2,15–18^. This is followed by a clustering of single-cells’ transcriptional profiles and/or inference of developmental trajectories^19,20^. Normally, the clusters or trajectory segments are validated and identified using expression of marker genes^11–14,17,21,22^. Another way to examine development trajectories more mechanistically is by inferring partial correlation networks (often referred to as regulatory networks) from the scRNA-seq data in question^11,17,22–24^. These standard approaches to scRNA-seq analysis outlined above help to deduce new insights from scRNA-seq. However, no method exists for the extraction of mechanistic patterns (i.e. gene modules) that explain cellular developmental programs over time. In bulk RNAseq data analysis, typically network enrichment technology is applied (e.g. KeyPathwayMiner^25^, GiGa^26^ or ActiveModules^27^). No such tool for scRNA-seq data exists, though. The main problem is the many missing values in typical scRNA-seq vs. bulk RNAseq data sets, as well as the necessity to analyse (pseudo)time-series data when comparing scRNA-seq trajectories (rather than typical case/control bulk RNAseq settings). To take full advantage of the potential of scRNA-seq data for extracting mechanistic patterns driving cell differentiation and cellular developmental programs, we have developed Scellnetor. The first (pseudo)temporal scRNA-seq network enrichment technique that can acquire a significantly more informative picture of the gene expression changes during cell development, as it identifies entire modules of interacting genes that are expressed similarly while the cells differentiate - or through pseudotime. To a certain extent, this can be addressed with existing tools by the identification of partial correlation networks underlying differentiation trajectories of single cells^28–30^. Despite being useful in specific scenarios, such (pseudo) gene regulatory networks are limited in their power to describe the interactome beyond transcription factors and do not grant a view on the full picture of the complexity of cellular developments. Currently, no tools exist to identify molecular subnetworks enriched with genes that are differently expressed in two distinct differentiation trajectories, thus unraveling how interacting genes “push” cellular development in certain directions as ensemble (i.e. mechanistic module). From a systems medicine point of view, as no tool for direct comparison between healthy differentiation trajectories and disease-associated development trajectories exists, it remains impossible to locate genes that in synergy, as mechanism, are responsible for disease progression. To fill this gap, we have invented Scellnetor: Single cell Network Profiler for Extraction of Systems Biology Patterns from scRNA-seq Trajectories. Essentially, Scellnetor is a clustering algorithm tailored to scRNA-seq data that enables the user to investigate pre-defined clusters or paths on low dimensional cell maps. The selected sets of cells will be sorted according to pseudotime and clustered using a hierarchical clustering algorithm that is constrained by the protein-protein interaction (PPI) network from BioGrid^31^ (or a user-chosen network). Scellnetor identifies connected subnetworks of genes that are either differently or similarly expressed in two selected sets of cells (Fig. 1). Such subnetworks will be referred to as gene modules throughout the paper. Scellnetor is, therefore, able to extract mechanisms that are fundamental cellular driver programs for differentiating between distinct development courses. Scellnetor is available as an intuitive and interactive online tool at https://exbio.wzw.tum.de/scellnetor.

**Figure 1.**
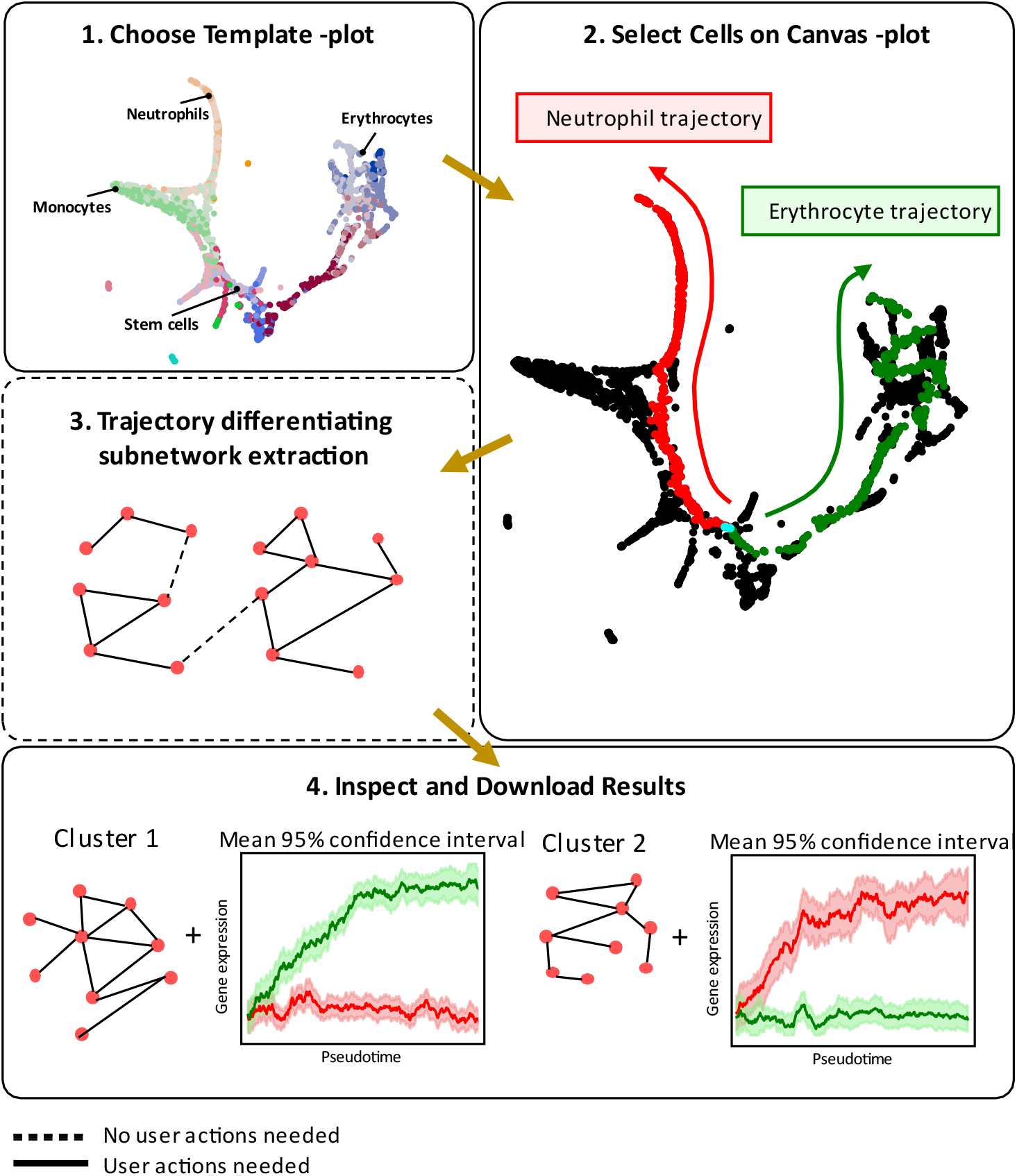
Workflow of Scellnetor. 1 Choosing desired Template-plot. **2** Selection of cells on Canvas-plot that represents differentiation trajectories. **3** Scellnetor performs a constrained hierarchical clustering based on expression data from the selected cells. **4** User can inspect and download the results from the Scellnetor clustering. Scellnetor outputs subnetworks of genes and mean gene expression with 95% confidence intervals.

## Results

### Differences between neutrophil and erythrocyte development trajectories

To validate the Scellnetor methodology, we used scRNA-seq data from ref. ^21^. In their article, Paul *et al.* analyzed gene expression patterns of mouse hematopoietic cells while they differentiated to progenies from a pool of progenitor cells: common myeloid progenitors (CMPs), granulocyte-macrophage progenitors (GMPs), and megakaryocyte-erythrocyte progenitors (MEPs). Using an

Expectation Maximization-based clustering approach, Paul *et al.* divided the single cells into 19 different groups. Finally, they constructed a detailed map of the dynamic transcriptional states within the myeloid progenitor populations. Using the single cells from the 19 clusters and their scRNA-seq gene expression data set (GSE72857) we constructed an ANNDATA object, created cell maps and computed pseudotime. Our ANNDATA object contained 2730 single cells that expressed 3451 genes (see Method section for more info). Our pseudotime calculation was based on progenitor clusters defined by Paul *et al.* (2015) (cluster 7-11 in Fig. S1) and coordinates from a cell map based on principles of force-directed graph drawing ^32^, which showed clear branching of the differentiated cells. Also, the plot showed clear co-localization of Paul *et al.* clusters that were highly similar. The “start cell” for the pseudotime computation^7^ was the cell closest to the average position of all relevant progenitor clusters (cluster 7-10 in Fig. S1, Fig S5). The cluster 11 was omitted, as it was dislocated from the remainder of the cells on the plot (Fig. S1). The connected area where Paul *et al.* clusters 7-10 are co-localized will be referred to as “stem cell area” (Fig S1, Fig S5).

We ran our Scellnetor algorithms to extract comparative systems biology profiles between 1.) the trajectory from stem cells towards differentiated neutrophils versus 2.) the trajectory from stem cells towards differentiated erythrocytes (Fig 2a). The drawn paths go through several of the Paul *et al.* (2015) defined clusters. Outside the stem cell area, the neutrophil path goes through clusters 15-17, where 16-17 are the neutrophil clusters. The erythrocyte path goes through cells in the stem cell area and the clusters 1-6, which all are erythrocyte clusters (Fig 2a, Fig S1). We identified seven clusters with a minimum size of five genes using pseudotime as sorting metric. The distance metric was Euclidean, the linkage type was complete and the size of the moving average was 20.

**Figure 2.**
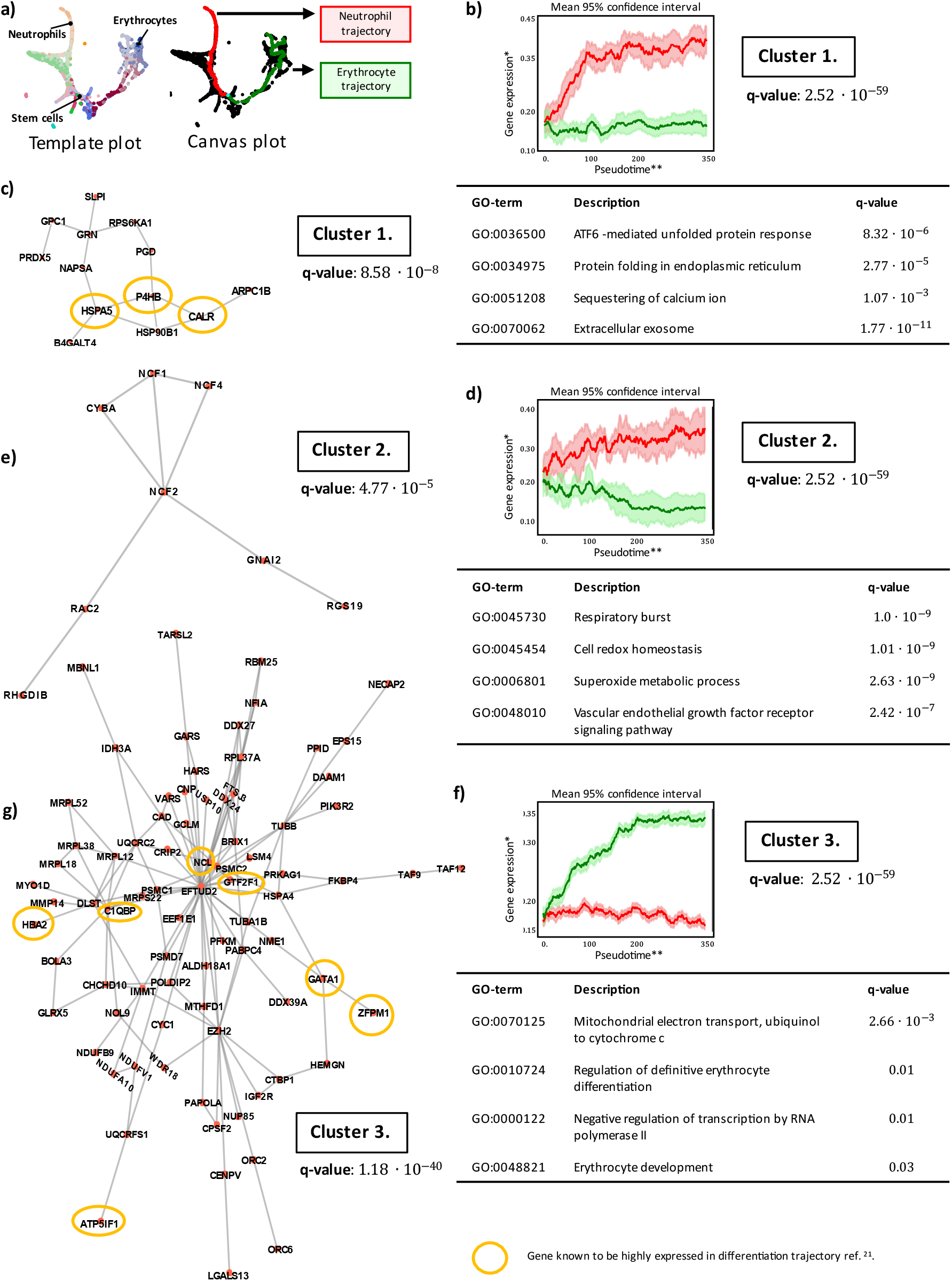
Comparison of neutrophil and erythrocyte differentiation trajectories. **a** Chosen Template-plot (left) and selected cells on Canvas-plot (right). The selected cells form paths representing trajectories from stem cells towards differentiated neutrophils and from stem cells towards differentiated erythrocytes, respectively. **b, d, f** Plots of mean pseudo-timelines and 95% confidence interval of cluster 1, 2 and 3. The mean pseudo-timelines from both trajectories of each cluster were compared using Wilcoxon signed-rank tests. Resulting q-values can be seen below every plot (p-values are adjusted using Benjamini-Hochberg Procedure). Below every plot are tables with four statistically significant GO-terms linked to the clusters. **c, e, g** Gene modules corresponding to cluster 1, 2 and 3. The encircled genes are known to be highly expressed in the differentiation trajectory that on average has the highest expression in **b, d, f** ref. ^21^. The associated q-values are based on Mann-Whitney U tests and indicate that these clusters are statistically significantly different from randomly generated clusters (p-values are adjusted using Benjamini-Hochberg Procedure). *Gene expression = Normalized moving-average-modified gene expression, **Pseudotime = Moving-average-modified number of single cells.

### Pseudo-timelines and validation

In Figure 2, we show three of the clusters. Figure 2b, d, and f show plots of the mean and 95% confidence intervals of the smoothed average expression in the three clusters (see supplementary method section). Below the plots are tables with GO-terms linked to each cluster (see next section). The red lines on the plots indicate the expression values of the clustered genes when following the path from stem cells to neutrophils while the green lines correspond to expression on the path from stem cell to erythrocytes. One can see how the modules’ genes’ expression differs over pseudotime between the neutrophil and erythrocyte trajectories. Using Wilcoxon signed-rank test, we show that the average expression over time of the clustered genes from the two distinct differentiation paths are statistically significantly different (q-value 2.52o 10^-59^ for all clusters).

### Gene expression of cells in Scellentor paths is consistent with gene expression in predefined clusters

In Figure 2a, the neutrophil path passes through stem cells and cells from Paul *et al.* clusters 15-17, whereas the erythrocyte path goes through stem cells and cells from Paul *et al.* clusters 1-6. The paths contain subsets of these clusters, so it was of interest to check if the differences in expression between the paths were consistent with the differences in expression between the corresponding clusters (e.g. clusters 15-17 vs. clusters 1-6 according to ref. ^21^). We extracted all genes from the Scellnetor clusters (Fig. 2, Fig S2) and created two cell sets; one containing all cells from the Paul *et al.* clusters 15-17 and one containing all cells from the Paul *et al.* clusters 1-6. Based on the two sets, we used our initial count matrix (see method section) to compute the average expression of the genes in the Scellnetor clusters. We plotted the averaged gene expression values and compared the resulting distributions using Wilcoxon signed-rank test (Fig. S3). The same was done for the subsets of the above Paul *et al.* clusters that were included in the paths shown in Figure 2a (Fig. S4). For these calculations, we used the preprocessed expression matrix from our ANNDATA object (see method section). We found that the cells in the Scellnetor paths (Fig. 2a), expressed genes in a manner that was consistent with the gene expression of the cells in the relevant Paul *et al.* clusters.

### Scellnetor clusters point towards erythropoiesis

The clustered genes as connected subnetworks can be seen on Figure 2c, e, g. The q-values next to every subgraph are based on Mann-Whitney *U* tests and indicate that these clusters are statistically significantly different from randomly generated clusters (q-values for cluster 1, 2, 3 are 8.58o 10^-8^, 4.77o 10^-5^, 1.18o 10^-40^, respectively). The genes *P4HB, CALR* and *HSPA5* in cluster 1 (Fig. 2c) produce surface markers that are expressed at high levels in neutrophils. The genes *GATA1, ZFPM1* and *GTF2F1* in cluster 3 (Fig. 2g) code for transcription factors that are upregulated in erythrocyte differentiating cells. The genes *NCL, HBA2, C1QBP* and *ATP5IF1* are all known marker genes associated with the erythrocyte lineage (Fig. 2g). Additional transcription factors upregulated in erythropoiesis are *GFI1B, LMO2* and *CBFA2T3*, which were found in Scellentor cluster 7 (Figure S2h)^21^. The genes in the Scellentor cluster 3 (Fig. 2g) are more highly expressed on average in the cells located in the erythrocyte trajectory. This is corroborated by higher expression of *HBA2*, which codes for a subunit of hemoglobin^33^ and *ATP5IF1*, which codes for a mitochondrial ATPase inhibitor that is involved in the in the synthesis of hemoglobin^34^.

### Gene Set Enrichment Analysis Results

Scellnetor can automatically conduct gene set enrichment analysis when gene modules have been identified. To demonstrate this functionality, we display four biologically meaningful statistically significant biological process GO-terms associated with each cluster in the tables in Figure 2b, d, f. To see all GO-terms associated with the clusters, see supplementary file 1. Genes involved in “ATF6-mediated unfolded protein response”, “protein folding in endoplasmic reticulum” and “sequestering of calcium ion” have been found highly expressed in cluster 1 (Fig. 2b). The first two GO-terms describe events that have to do with protein folding and endoplasmic reticulum (ER) stress. It has been shown that ER stress is decreased during neutrophil differentiation^35^. Scellnetor uncovered that *CALR* is highly expressed in the neutrophil trajectory compared to the erythrocyte trajectory. *CALR* code for a multifunctional protein called calreticulin, which counteracts ER stress and is involved in correct maintenance of calcium ions in the ER^36–40^. The gene *P4HB* (Fig 2e) have also been found to downregulate ER stress^41^. Eleven out of 13 genes in cluster 1 (Fig. 2b) (except for *B4GALT4* and *RPS6KA1)* are associated with the GO-term “extracellular exosome”. Releasing exosomes is an important part of neutrophil signaling^42^. The GO-terms “respiratory burst”, “cell redox homeostasis” and “superoxide metabolic process” from cluster 2 (Fig. 2d) all relate to well-known neutrophilic cellular processes. A distinctive feature of the inflammatory actions of neutrophils is the respiratory or oxidative burst, where large amounts of oxygen are consumed to produce superoxide^43^. Recent studies revealed that neutrophils might play a key role in angiogenesis, as they can store and synthesize molecules with known angiogenic activity (fourth GO-term, Fig. 2d)^44–47^. Cluster 2 does not contain any genes that were highlighted by ref. ^21^ as noteworthy for the neutrophilic differentiation course. Still, it has been described that the neutrophil cytosolic factor 1, 2, 4 (*NCF1, NCF2, NCF4)* interact with *CYBA* and *RAC2* at the membrane as subunits of the NOX2 complex. Interestingly, the inclusion of *RAC2* in this molecular assembly is specific for neutrophils^48–50^.

The GO-terms “regulation of definitive erythrocyte differentiation” and “erythrocyte development” from cluster 3 (Fig. 2f) indicate that the cells from the erythrocyte trajectory in fact do express genes associated with erythropoiesis. The GO-term “mitochondrial electron transport, ubiquinol to cytochrome c” (Fig. 2f) is interesting, because the proteins resulting from the genes linked to this term (*UQCRC2, UQCRFS1, CYC1*, Fig. 2g) have been found in high abundance in erythroid progenitor cells compared to hematopoietic stem cells^51^. Apparently, these genes are higher expressed in erythrocyte differentiating cells than in neutrophils differentiating cells. The GO-term “negative regulation of RNA polymerase II” fits well with genes involved in erythropoiesis, since mature erythrocytes lose their nuclei, which is where RNA polymerase II catalyze transcription of genes to pre-mRNA^52^.

### Scellnetor identifies trajectory-explaining gene modules for functional and dysfunctional exhausted CD8 T cells in chronic infection

To demonstrate that Scellnetor can be used for identifying mechanisms underlying disease progression, we reanalysed a dataset of pathways of CD8 T cell differentiation in a well-defined standard model of chronic infections^12^. In the original work, the authors investigated proliferation and differentiation of CD8 T-cell progenitors to effector cells in populations of dysfunctional and normal CD8 T-cells with and without CD4 T-cell help. They revealed that absence of CD4 T-cell reduces the number of terminally differentiated CD8 T-cells, but not the number of CD8 T-cell progenitors in exhausted T-cell populations. Thus, the progenitor cells can maintain their population size without CD4 T-cell help. They identified 5 clusters and labeled them based on previously established signature genes they; cluster 1 represented the critical stem-like progenitors, cluster 2 the functional and clusters 3, 4 and 5 the dysfunctional effector cells. In our study, we used the cells from these 5 clusters (GSE137007) to generate an ANNDATA object, calculate pseudotime and compute a diffusion map (see method section). We used their cluster annotations to color code the cells (Fig 3a).

**Figure 3.**
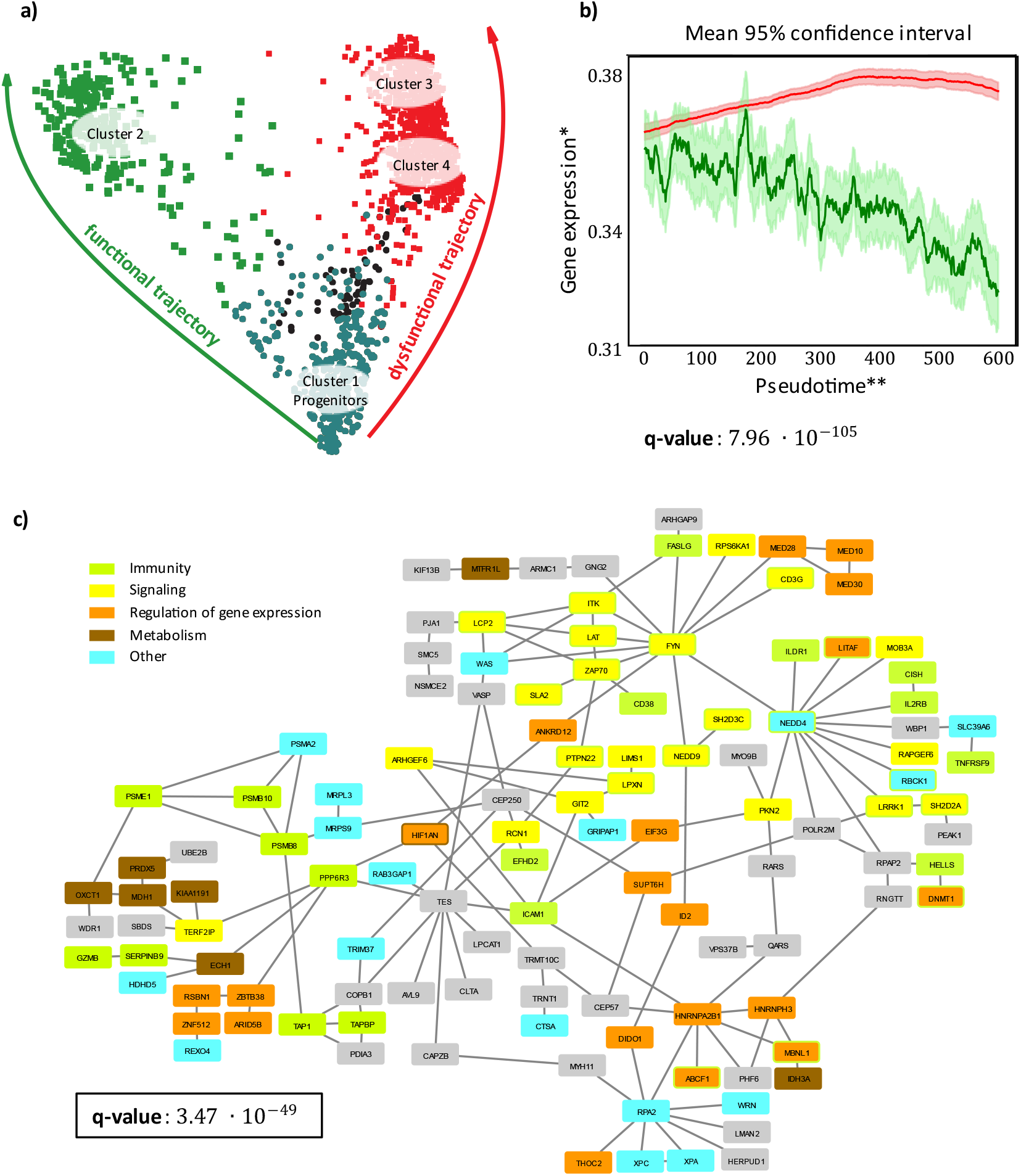
Differentiation of progenitor cells to dysfunctional CD8 T-cells. **a** Diffusion map of the single cells as they differentiate to functional and dysfunctional CD8 T-cells. On the left, in green, is the development trajectory of cells from progenitor cells (cluster 1 in ref. ^12^) to functional CD8 T-cell (cluster 2 in ref. ^12^). On the right, in red, is the development trajectory of cells from progenitor cells to dysfunctional CD8 T-cell (cluster 3, cluster 4 in ref. ^12^). **b** Mean gene expression and 95% confidence intervals of the selected cluster. The genes in the cluster shown in **c** are, on average, expressed at a higher level in the cells developing via the functional trajectory. The q-value below the plot is based on a Wilcoxon signed-rank test comparing the average gene expressions. **c** The module induces by the cluster of genes. The genes are color coded; green genes are involved in immune responses, yellow genes are involved in signaling, orange genes are involved in regulation of gene expression, brown genes are involved in metabolism, blue genes are involved in other processes and grey genes do no belong to any of the mentioned groups. The color coding and, thereby, grouping of genes is based on our subjective assessments of their general functions. The associated q-value is based on Mann-Whitney U test. It indicates that the cluster is statistically significantly different from randomly generated clusters. All p-values in this figure are adjusted using Benjamini-Hochberg Procedure. *Gene expression = Normalized moving-average-modified gene expression, **Pseudotime = Moving-average-modified number of single cells.

We defined two trajectories of differentiation starting from the progenitor subpopulation (cluster 1), one towards the functional effector cells (cluster 2) and one towards the dysfunctional effector cells (cluster 3 and 4). We excluded cluster 5 (Figure 3a, cells in black) from the analysis as it is conditions-pecific and it was only observed in the absence of CD4 help. Using Scellnetor, we unravelled a gene module with increased expression in cells progressing in the dysfunctional trajectory and decreased expression in those progressing in the functional trajectory (Fig. 3b, c). The complexity of the gene module is highlighted by the diversity of genes and their functions, including processes like regulation of gene expression (epigenetic, transcriptional and translational), signalling, immune defence, cell migration, cytoskeletal reorganization and genes involved in the T cell receptor signalling pathway (*CD3G, FYN, ZAP70, LAT, ITK, PTPN22, LCP2, SLA2, NEDD4, NEDD9, LRRK1, SH2D2A* and *SH2D3C)*, whose excessive stimulation is known to be one of the main drivers of T cell exhaustion^53^. The two most statistically significant GO-terms associated with this cluster are “T-cell receptor signaling pathway” and “T-cell activation” (see supplementary file 2). TCR signalling is connected to two therapeutically interesting receptors involved in regulation of the T cell response – ILRD1 and TNFRSF9, which could potentially be used for its modulation. ILDR1 function as regulator of T cells response in chronic infection and cancer is intriguing, especially taken into account that ILRD2 was recently described as a negative regulator of the T cells^54^. However, up to now it has never been linked to controlling CD8 T cell function neither in chronic infection nor in cancer. TNFRSF9 (coding 4-1BB) is a positive regulator of T cell effector function and survival, and its stimulation has already been reported to ameliorate T cell exhaustion^55^. The mechanistic gene module also includes diverse regulators of NF-κB signaling (PPP6R3, TERF2IP and EFHD2), which is critical for T cell survival and cytokine production^56^. The negative regulator EFHD2 is of particular interest, as it is necessary for PD-1 mediated inhibition of proliferation and cytokine secretion in dysfunctional CD8 T cells^57^. As previously reported, the transition from functional to dysfunctional CD8 T cells is associated with metabolic reprogramming marked by a switch from glycolysis to oxidative phosphorylation (OXPHOS) as a main pathway for generation of adenosine triphosphate (ATP). In line with this, our dysfunctional gene module includes multiple genes involved in fatty-acid beta oxidation and OXPHOS (MDH1, KIAA1191, IDH3A, ECH1, OXCT1 and PRDX5) and putative regulators of this metabolic adaptation (HIF1AN, ARID5B and MTRF1L). Interestingly, HIF1AN is an inhibitor of HIF1α, which is known to trigger the expression of genes promoting the use of glycolysis over mitochondrial oxidative phosphorylation as main energy generating pathway^58–60^. Moreover, HIFs activity has been shown to enhance the effector CD8 T cell response and influence the expression of pivotal transcription, effector and co-stimulatory molecules in chronic infection^61^. Altogether, this demonstrates that the gene modules extracted by Scellnetor are functionally related mechanisms and precisely reflect the opposing nature of progenitor differentiation towards functional or dysfunctional CD8 T cells. Thus, Scellnetor is a promising systems medicine hypothesis generator identifying molecular sub-networks mechanistically driving dynamic differentiation of complex cell populations.

## Methods

Scellnetor is a novel algorithmic approach implemented into a responsive webtool that clusters scRNA-seq data (Fig. 4). It starts with a molecular interaction (usually: PPI) network, as well as a user-generated cell map from user-given scRNA-seq data. The coordinates of the cells are extracted and plotted on a canvas, where users may interactively select two trajectories or two sets of pre-defined clusters. Trajectories are selected by drawing paths on the canvas-plot, which are then connected by a greedy path-finding and smoothing algorithm. Clusters can be compared using subnetwork enrichment for differentiating expression module extraction. In contrast, trajectory cells are sorted along pseudotime^7^ to extract differentially expressed temporal subnetwork modules. Note that the webtool also allows identifying similarly (not differentially) expressed subnetwork modules, both for clusters and trajectories. Essentially, our network enrichment algorithm relies on computing a hyper-similarity matrix (genes vs. genes) that reflects the distance of pairs of genes in the network and their (dis)similarity of expression between the two selected clusters or over the two selected trajectories. This matrix is then clustered hierarchically and the induced subnetwork of emerging gene clusters are reported as candidate subnetwork modules. Scellentor’s output are 1) the corresponding subnetwork modules and the corresponding gene sets together with 2) plots of the mean expression development (over pseudo-time) of the genes in the modules including the 95% confidence intervals, and 3) statistically significantly module-associated GO-terms. For a more detailed description of the Scellnetor methodology see supplementary method section and Fig S7.

**Figure 4.**
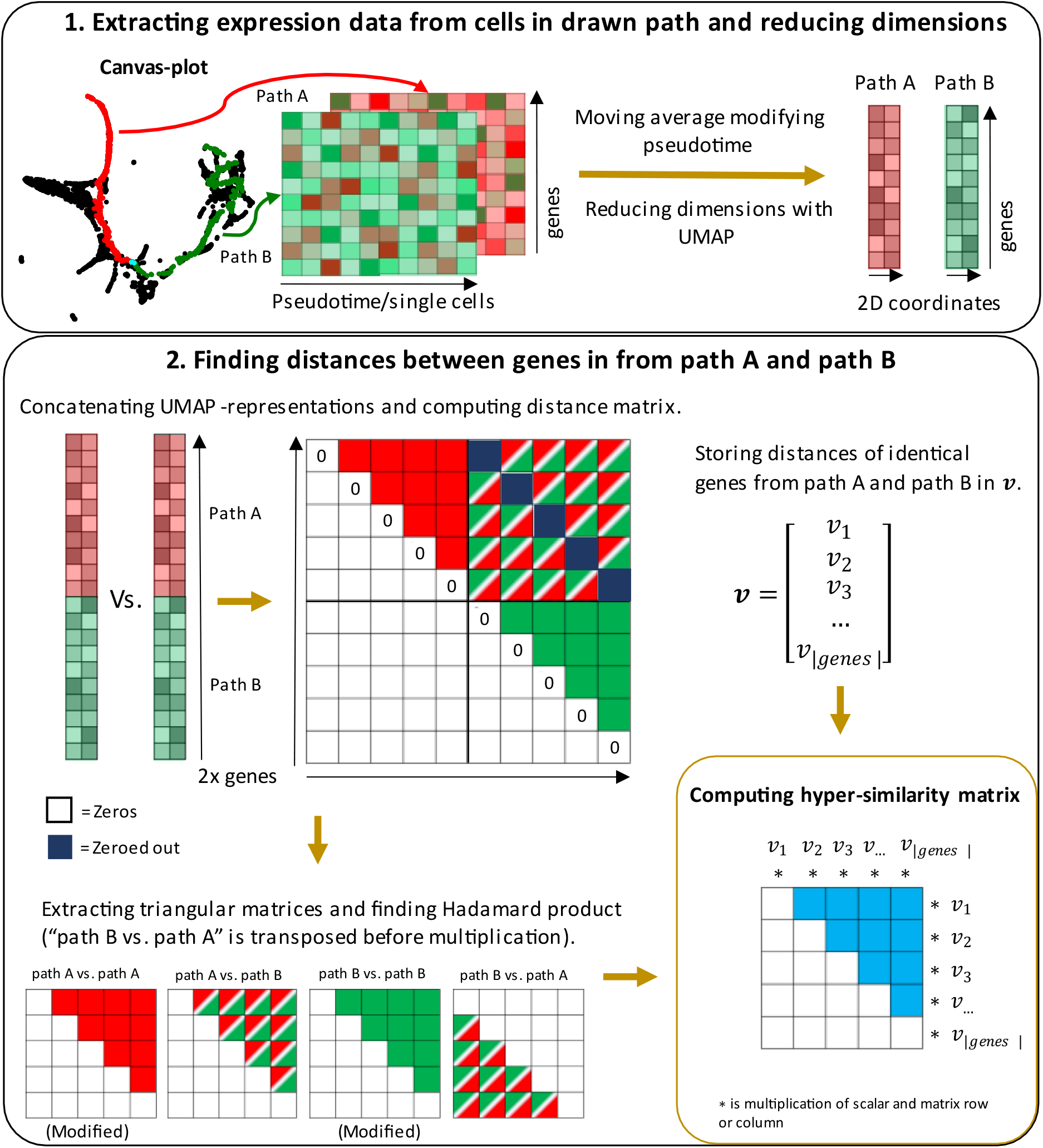
Computation of hyper-similarity matrix for gene module discovery. **1** Two paths are drawn on the canvas-plot, path A and path B. Messenger-RNA expression data are extracted from selected cells and used to create expression matrices - one for each drawn path. To ease the understanding of the method, all data are derived from path A or path B, will be labeled as path A or path B in the figure. Path A and path B are converted into expression matrices, where rows are genes and columns are cells (pseudo-timepoints). Each row of the expression matrices is smoothed using a moving average function. Afterwards, the smoothed expression matrices are dimensionality reduced with UMAP, such that every pseudo-timeline of genes in the two sets are converted to 2D coordinates in the Euclidean space. **2** The 2D coordinates from path A and path B are concatenated along their first axes. The concatenated coordinate sets are used to produce a distance matrix, which is normalized by division of its max value. The diagonal, the lower triangular matrix and the diagonal of the upper right quadrant of the distance matrix are zeroed out. The distances between genes with identical IDs, but from different sets, are computed and stored in the vector, *𝒗* (top, right). Triangular matrices of the distance matrix’ quadrat I, II and IV (as in cartesian coordinate system) are extracted. Matrices “path A vs. path A” and “path B vs. path B” are modified such that genes with the shortest distances have the highest values and vice versa. The matrix “path B vs. path A set” is trans posed and the Hadamard product of all triangular matrices is computed (bottom, left). This produces the hyper-similarity matrix. The rows and columns of the hyper-similarity matrix are weighted by the values of *𝒗* (bottom, right).

## Discussion

Despite the wealth of different tools designated for the analysis of scRNA-seq raw data, *de novo* identification of interacting subnetworks expressed in a similar fashion across pseudotime has been neglected so far. With Scellnetor, we present the first algorithm (implemented as intuitive webtool and stand-alone software) for directly identifying modules of connected genes with similar expression patterns along pseudo-time. Furthermore, Scellnetor allows for the comparison of two user-selected trajectories and finding gene modules that are either similarly of differently expressed through pseudotime. It uncovers interacting genes that are involved in the cellular differentiation.

To showcase this, we used Scellnetor to cluster scRNA-seq data from a hematopoiesis study by ref. ^21^. We compared a developmental trajectory from stem cells towards differentiated neutrophils with a trajectory from stem cells towards differentiated erythrocytes. We uncovered statistically significant gene modules that are involved in differentiation of neutrophils and erythrocytes, respectively. The identified modules contained genes that previously have been associated with the two distinct differentiation courses and genes with GO-terms that were closely connected to the analyzed cellular developments (Fig. 2), demonstrating the biological relevance of the discovered modules.

We also applied Scellnetor to examine diseases progression by analyzing scRNA-seq data from ref. ^12^. We compared two trajectories starting from the progenitor subpopulation (cluster 1 in ref. ^12^); one towards the functional effector cells (cluster 2 in ref. ^12^) and one towards the dysfunctional effector cells (cluster 3 and 4 in ref. ^12^) (Fig. 3). Scellnetor found several statistically significant clusters, of which we highlight one module consisting of genes that are, on average, significantly higher expressed in cells developing in the dysfunctional trajectory than in cells developing along the path towards “healthy” effector cells (Fig. 3). This module contains many genes that have already been associated with dysfunctional CD8 T-cell development but also new genes that offer new insights into the mechanistic foundation of CD8 T-cell development.

To sum up, Scellnetor employs a novel network-constrained hierarchical agglomerative clustering algorithm, which enables the comparison of two sets of single cells taking scRNA-seq data and a molecular interaction network as input. It is the first single cell network enrichment method allowing the extraction of systems biology patterns from scRNA-seq trajectories. In two real-world data sets we demonstrated its potential to extract disease progression mechanisms and cellular programs driving cell differentiation.

## Software availability

Scellnetor is freely available as an online tool at https://exbio.wzw.tum.de/scellnetor/ and can be downloaded as standalone program from GitLab (https://gitlab.com/AlexTheKing/scellnetor_docker_public).

## Data availability

The scRNA-seq data used for the Scellentor hematopoiesis clustering is from GEO (GSE72857). The scRNA-seq used for the clustering of exhausted CD8 T-cells in chronic infections is also from GEO (GSE137007). All Scellnetor results can be downloaded from GitLab (https://gitlab.com/AlexTheKing/scellnetor_docker_public).

## Code availability

On GitLab (https://gitlab.com/AlexTheKing/scellnetor_docker_public) one may find the analysis scripts and data representations used for the preparation of the manuscript.

## Supplementary data

- Supplementary method
- Supplementary figures
- Supplementary figure legends
- supplementary_file_1.xlsx
- supplementary_file_2.xlsx

## Acknowledgements

JB and AG received funding from JB’s VILLUM Young Investigator Grant nr. 13154. The work of JB and TK was further funded by H2020 project RepoTrial (nr. 777111). The work of RR and JB has partially been funded by H2020 project FeatureCloud (nr. 826078). JB and TK grateful for financial support from BMBF project Sys_Care. MO is grateful for financial support of the Collaborative Research Center SFB924.

## Author contribution statements

AG, JB and RR conceived the idea of the Scellnetor pipeline. AG developed and implemented the clustering algorithm of the Scellnetor tool. AG developed and implemented all basic backend functionalities of the webtool. AG, JL and MO further developed the webtool and significantly contributed to debugging. KK, TK, and DZ tested the webtool, provided critical feedback, and together with AG used Scellnetor to generate the biomedical result presented in the manuscript. All authors equally contributed to writing and improving the paper.

