## Supplementary method for "Comparative single-cell trajectory network enrichment identifies pseudo-temporal systems biology patterns in hematopoiesis and CD8 T-cell development"

### Supplementary Method Section

#### Introduction to Scellnetor

Scellnetor is the first network-constraint time-series clustering algorithm implemented as interactive webtool to identify modules of genes connected in a molecular interaction network that show differentiating temporal expression patterns. Scellnetor allows unraveling network modules driving, for instance, cell differentiation or disease progression on a system biological level. In addition, the mechanisms identified by Scellentor can be utilized for developing mechanistically oriented drugs that work on subnetwork-defined disease endophenotypes. Scellnetor is available as an intuitive and interactive online tool at https://exbio.wzw.tum.de/scellnetor and for download at GitLab (https://gitlab.com/AlexTheKing/scellnetor_docker_public).

Scellnetor requires  Scanpy-generated ANNDATA[1] objects as raw data input and scanpy-generated plots as template-plots. The user can interactively select cells of interest for the network module identification (Fig. S6). In order to form a time-series, the cells are sorted according to pseudotime, since it contains information about differentiation statuses of cells[2] (sorting according to an alternative key available in the ANNDATA object is supported as well). One of Scellnetor's key features is the comparison  of two distinct sets of single cells, which are either two paths through the template-plot or two sets of clusters, respectively. Scellnetor will compute a hyper-similarity matrix when comparing two sets. The hyper-similarity matrix contains information on how the genes are expressed compared to one another in the individual sets and compared to the genes in the set they are compared against. In case the user does not want to compare two sets of cells but look for the network module showing the most consistent expression profile, a standard distance matrix is calculated.

Scellentor uses a constrained hierarchical agglomerative clustering algorithm for extracting subnetworks of genes. Per default, the clustering is constrained by the interactions of the human PPI network from BioGrid[3], which makes it possible for Scellnetor to output clusters as connected subnetworks of genes. When comparing two paths or sets of clusters, the user can choose between searching for clusters of genes that are either differently or similarly expressed in the two sets. In addition to the connected subnetworks, Scellnetor outputs TSV files with statistically significant GO-terms for each cluster and plots of the mean and 95% confidence intervals of the expression patterns of the genes in the clusters.

The focus of this section will be on the functionalities associated with path-drawing and calculation of the hyper-similarity matrix, since these are the main novelties of the Scellnetor methodology. To quickly find out what a hyper-similarity means, see subsection “Summary - what do hyper-similarity values mean?”. All examples in this section will assume that the user has drawn paths on a canvas-plot. From the sub-section “Extraction of cells and smoothing of timelines” till the end of this section, the data processing for sets of clusters and user-drawn path is the same.

#### Extraction and conversion of genes from uploaded file

As mentioned before, Scellnetor constrains the clustering by the interactions of the Biogrid network. Nevertheless, the user can also utilize any other network provided in a tab separated edge list. The only restriction is the required usage of human Entrez IDs as node identifiers. The same restriction applies to the genes of uploaded H5AD files, which also are required to be given as EntrezID (or as identifiers that can be converted to Entrez IDs) and are additionally available as nodes in the network. All IDs without a corresponding network node will be discarded from further analysis. Scellnetor automatically recognizes human Entrez IDs, human gene symbols, human Ensemble stable IDs and mouse gene symbols. Due to overlapping gene symbol usage and identical human-mouse gene orthology of certain genes, some human and mouse gene symbols are mapped to the same human Entrez IDs. In these cases, Scellentor will automatically represent these Entrez IDs by the first instance of the genes in question that Scellnetor meets in its search. We recommend that users integrate a list with either human Entrez IDs or Ensemble stable IDs as an ANNDATA.var column, since this will preserve the highest number of genes that can be used for the clustering.

#### Selecting clusters on the canvas-plot

Users select cells on a canvas-plot, and the cluster to which the cell belongs is colored in accordance with the chosen set-color (Fig. S6). Switching between colors and selecting cells in different clusters makes it possible to create  two distinct sets of clusters that can be compared.

#### Drawing paths on canvas-plot

When drawing paths with Scellnetor, users select cells on the canvas-plot, which will be automatically connected to a path. The order in which the cells are selected directs the construction of the path through the plot. Let $Z$ be the set of all cells and $X = \{x_1, x_2, x_3 \dots x_n\}$ the set of user-selected cells. Then the path is created by forming a path between $x_1$ and $x_2$ followed by a path between $x_2$ and $x_3$ and so forth.. A path between cell $x_i$ to cell $x_{i+1}$ is generated by finding the cell, $x_{ia}$, that is closest to $x_i$ and closer to $x_{i+1}$than $x_i$ is. This is followed by finding the cell, $x_{ib}$, that is closest to $x_{ia}$ and closer to $x_{i+1}$than $x_{ia}$ is. This is repeated until a sub-path between $x_i$ and $x_{i+1}$ has been drawn. This is done for all possible pairs of $x_i$ and $x_{i+1}$ in $X$. The result is a minimal greedy path, which is the path connecting the cells in $X$ with the fewest number of cells possible when following the rules of the greedy path algorithm. The cells in the resulting path are stored in the set, $X_{path}$.

#### Expanding trajectories

The paths can be expanded (i.e., made thicker to include more cells) by including a user-defined percentage of cells that neighbor cells in the minimal greedy path. The user can expand the paths to contain $pct$ percentage of the overall dataset. This is achieved by greedily adding the closet cells to cells which are already in $X_{path}$ until the desired number of cells is reached. Euclidean distance is used as distance metric and $X_{expand} = X_{path}$ when $i = 0$ for $X_{path} = \{x_1, x_2, x_3 \dots x_N\}$ where $N$ is the number of cells in the minimal greedy path.

#### Finding similar-sized paths

For optimal comparisons of paths, it is recommended that users make paths that contain the same number of cells (see Extraction of cells and smoothing of timelines for an explanation). Scellnetor has integrated functionalities that allow users to even out the sizes of their drawn trajectories. If two paths have been drawn and one has been expanded by inclusion of the $pct$ percent closest points, and the other is in its minimal greedy path form, then let $X_{expanded}$ be the set containing the cells of the expanded trajectory and $Y_{path}$ be the set containing the cells of the path, which is in its minimal greedy path form. To even out the sizes of $X_{expanded}$ and $Y_{path}$, $Y_{path}$ is first expanded by the inclusion of the $pct$ percent closest cells for every $y_i \in Y_{path}$ (see Expanding trajectories). The result is the set $Y_{expanded}$. If $|Y_{expanded}| = |X_{expanded}|$, then the objective has been reached. If $|Y_{expanded}| < |X_{expanded}|$, a cell from $Y_{expanded}$ is chosen at random and its closest neighbor that is not already in $Y_{expanded}$, is added to the set. This is repeated until $|Y_{expanded}| = |X_{expanded}|$. Adding cells like this will promote a uniform expansion of the trajectory. If $|Y_{expanded}| > |X_{expanded}|$, distances from the cells in $Y_{path}$ to cells in $Y_{expanded} - Y_{path}$ are calculated. And the cells in $Y_{expanded} - Y_{path}$ that have the average longest distance to all cells in $Y_{path}$ are removed until $|Y_{expanded}| = |X_{expanded}|$. Users can even-out their paths even if both paths have been expanded or none of them has. However, it is not possible to get a path smaller than the path's minimal greedy path form.

#### Extraction of cells and smoothing of timelines

A hyper-similarity matrix is only calculated when two sets are compared. From this subsection onwards till the subsection "Generation of results", it will be assumed that a user has drawn two paths A and B. Path A and path B are converted into matrices $\boldsymbol{A}$ and $\boldsymbol{B}$, respectively. Matrix $\boldsymbol{A}$ has size $g \times m$ and $\boldsymbol{B}$ has size $g \times l$, where $g$ is the number of genes, and $m$ and $l$ are the number of cells in path A and path B, respectively. Per default, a moving average function is applied on the rows of $\boldsymbol{A}$ and $\boldsymbol{B}$, which results in new matrices called $\boldsymbol{A}_{mean}$ and $\boldsymbol{B}_{mean}$ with sizes $g \times m'$ and $g \times l'$, respectively. The element in row $i$ and column $j$ of $\boldsymbol{A}_{mean}$ and $\boldsymbol{B}_{mean}$ are found using the following moving average formula:

$$A_{mean_{i,j}} = \frac{1}{a}\sum_{z=0}^{a-1} A_{i,j+z} \tag{1}$$

where $a$ is the user-defined size of the moving average and $A_{i,j}$ is the element in row $i$ and column $j$ of matrix $\boldsymbol{A}$. Formula (2) is applied on every row of $\boldsymbol{A}$ and $\boldsymbol{B}$ and every $j$-th column in the range $\{j \in \mathbb{N} \mid 0 \leq j \leq m - a + 1\}$ for $\boldsymbol{A}$ and every $j$-th column in the range $\{j \in \mathbb{N} \mid 0 \leq j \leq l - a + 1\}$ for $\boldsymbol{B}$. In cases where the two paths are of different lengths, the moving average window of the longer path is adjusted such that both paths contain the same number of smoothed values. This means, if a path contains a much higher number of cells than the path it is compared to, the moving average can "flatline" the gene expression patterns of the larger set. Therefore, it is ideal if the objects to be compared have identical sizes.

#### UMAP of gene expression of single cells

Before the gene expression patterns in the two sets are compared, the smoothed data in $\boldsymbol{A}_{mean}$ and $\boldsymbol{B}_{mean}$ are dimensionality reduced. UMAP[4] is applied for the dimensionality reduction given one of the four user-chosen distance metrics: Euclidean distance, Manhattan distance, Minkowski distance or Correlation. Scellnetor sets the UMAP parameters *n_neighbors*=30, *min_dist*=0.0 and *random_state*=42. The remaining UMAP parameters are set to their default values. $\boldsymbol{A}_{mean}$ and $\boldsymbol{B}_{mean}$ are concatenated before transformed by UMAP, as the gene expression patterns from the two matrices thereby will be high-dimensional coordinates on the same manifold structure. This will provide more meaningful inter-coordinate distances of the dimensionality reduced data. UMAP outputs a different coordinate landscape depending on the order of the concatenation. For example, the distances between coordinates resulting from a UMAP-embedding of the concatenation of $\boldsymbol{A}_{mean}$ and $\boldsymbol{B}_{mean}$ are similar, but slightly different from the distances between coordinates resulting from a UMAP-embedding of the concatenation of $\boldsymbol{B}_{mean}$ and $\boldsymbol{A}_{mean}$.

So, to retain determinism of the Scellnetor clustering, a dimensionality reduction is conducted on both the concatenation of $\boldsymbol{A}_{mean}$ and $\boldsymbol{B}_{mean}$, and the concatenation of $\boldsymbol{B}_{mean}$ and $\boldsymbol{A}_{mean}$. The resulting dimensionality-reduced matrices, $\boldsymbol{C}_{small_{AB}}$ and $\boldsymbol{C}_{small_{BA}}$, both have the size $2g \times 2$, where $g$ is the number of genes in $\boldsymbol{A}_{mean}$ and $\boldsymbol{B}_{mean}$. $\boldsymbol{C}_{small_{BA}}$ is redefined as the concatenation of $\boldsymbol{C}_{small_{BA}}[g:]$ on top of $\boldsymbol{C}_{small_{BA}}[:g]$, as the order of the genes in the two coordinate sets should be identical for the further processing. $\boldsymbol{C}_{small_{BA}}[g:]$ is the slice of $\boldsymbol{C}_{small_{BA}}$ that contains the last $g$ rows and $\boldsymbol{C}_{small_{BA}}[:g]$ is the slice of $\boldsymbol{C}_{small_{BA}}$ that contains the first $g$ rows.

#### Hyper-similarity matrix

When comparing the two paths, path A and path B, a hyper-similarity matrix is computed. $\boldsymbol{C}_{small_{AB}}[:g]$ contains the moving average-modified and dimensionality reduced gene expression patterns from path A, and $\boldsymbol{C}_{small_{AB}}[g:]$ contains the moving average-modified and dimensionality reduced gene expression patterns from path B. The same holds true for $\boldsymbol{C}_{small_{BA}}[:g]$ and $\boldsymbol{C}_{small_{BA}}[g:]$, respectively. Again, $g$ is the number of genes in $\boldsymbol{A}_{mean}$ and $\boldsymbol{B}_{mean}$. A $\boldsymbol{C}_{small_{AB}}$ vs. $\boldsymbol{C}_{small_{AB}}$ distance matrix, $\boldsymbol{D}_{small_{AB}}$, and a $\boldsymbol{C}_{small_{BA}}$ vs. $\boldsymbol{C}_{small_{BA}}$ distance matrix, $\boldsymbol{D}_{small_{BA}}$, is calculated using Euclidean distance as metric. Again, to retain determinism of the Scellnetor clustering approach, a distance matrix, $\boldsymbol{D}_{small}$, is found by

$$\boldsymbol{D}_{small} = \frac{\boldsymbol{D}_{small_{AB}} \oplus \boldsymbol{D}_{small_{BA}}}{2}, \tag{2}$$

where the values of $\boldsymbol{D}_{small}$ are normalized by

$$\boldsymbol{D}'_{small} = \frac{\boldsymbol{D}_{small}}{max(\boldsymbol{D}_{small})}. \tag{3}$$

In the matrix $\boldsymbol{D}_{small}$ in our example (Fig S7), the upper left quadrant corresponds to a path A vs. path A distance matrix and the lower right quadrant corresponds to a path B vs. path B distance matrix. The upper triangle of the upper right quadrant corresponds to the upper triangle of a path A vs. path B distance matrix and the lower triangle of the upper right quadrant corresponds to the lower triangle of a path B vs. path A distance matrix.

The diagonal of $\boldsymbol{D}_{small}$ is zeroed out. The diagonal of the upper right quadrant of $\boldsymbol{D}_{small}$ is zeroed out as well. the latter will be explained in the subsection "Weighting values of the hyper-similarity matrix" below. The upper triangular matrix of the upper left quadrant is defined as $\boldsymbol{D}_{AA}$, the upper triangular matrix of the lower right quadrant is defined as $\boldsymbol{D}_{BB}$, the upper triangular matrix of the upper right quadrant is defined as $\boldsymbol{D}_{AB}$ and the lower triangular matrix of the upper right quadrant is defined as $\boldsymbol{D}_{BA}$. The values in $\boldsymbol{D}_{AA}$ and $\boldsymbol{D}_{BB}$ are reversed by

$$\boldsymbol{D}'_{XX} = \left|\left(\boldsymbol{D}_{XX} - (1 - 10^{-6})\right)\right| \tag{4}$$

where $\boldsymbol{D}_{XX}$ is the matrix and $10^{-6}$ is a small value subtracted from 1 to set the values of the matrix in the range ]0; 1]. This way, no values are zeroed out after reversion of the distance-values and the relative distance-differences remain unchanged. If the user wants clusters of connected genes that have similar expression patterns in path A and path B, respectively, and similar expression patterns when comparing path A and path B, then $\boldsymbol{D}_{AB}$ and $\boldsymbol{D}_{BA}$ are updated using (9) (*cluster type 1*).

The matrices $\boldsymbol{D}_{AB}$ and $\boldsymbol{D}_{BA}$ remain as initially defined, if the resulting clusters should be connected genes that have similar expression patterns in path A and path B, respectively, and dissimilar expression patterns when comparing path A against path B (*cluster type 2*). To compress the distances from $\boldsymbol{D}_{AA}$, $\boldsymbol{D}_{BB}$, $\boldsymbol{D}_{AB}$ and $\boldsymbol{D}_{BA}$ into a single "pre-hyper-similarity matrix", $\boldsymbol{D}$, the four matrices are element-wise multiplied as follows

$$\boldsymbol{D} = \boldsymbol{D}_{AA} \odot \boldsymbol{D}_{BB} \odot \boldsymbol{D}_{AB} \odot \boldsymbol{D}_{BA}^{\top} \tag{5}$$

where $\odot$ it the Hadamard product. The diagonal and the lower triangular matrix of $D$ are zeroed out. Now, the matrix $\boldsymbol{D}$ only needs a few processing steps before it is ready for the clustering as a hyper-similarity matrix. The matrices $\boldsymbol{D}_{AA}$, $\boldsymbol{D}_{BB}$, $\boldsymbol{D}_{AB}$ and $\boldsymbol{D}_{BA}$ corresponds to the matrices $\boldsymbol{D}_{rr}$, $\boldsymbol{D}_{gg}$, $\boldsymbol{D}_{rg}$ and $\boldsymbol{D}_{gr}$, respectively, on Figure S7.

#### Weighting values of the hyper-similarity matrix

In $\boldsymbol{D}$, the values depend on the distances between the gene expression patterns from the cells in path A and on the distances between the gene expression patterns from the cells in path B. The matrices $\boldsymbol{A}_{mean}$ and $\boldsymbol{B}_{mean}$, derived from the two paths, contain the moving average modified expression values of the same genes in the same order, but measured as the cells in the two sets follow different differentiation trajectories. The matrix element at $\boldsymbol{D}_{i,j}$ depends on the expression values of gene $i$ in $\boldsymbol{A}_{mean}$ and $\boldsymbol{B}_{mean}$, respectively, and on the expression values of gene $j$ in $\boldsymbol{A}_{mean}$ and $\boldsymbol{B}_{mean}$, respectively. The matrix element at $D_{i,j}$ should also depend on the distance of gene $i$ in $\boldsymbol{A}_{mean}$ vs.

gene $i$ in $\boldsymbol{B}_{mean}$ and gene $j$ in $\boldsymbol{A}_{mean}$ vs. gene $j$ in $\boldsymbol{B}_{mean}$. Until now, these values have been zeroed out and ignored, but they will be used to weigh the matrix $\boldsymbol{D}$ in the following steps (Fig. S7).

For example, if a user applies Euclidean distance as metric and wants to find clusters of *cluster type 1* then the value in row $i$ and column $j$ of $\boldsymbol{D}$, should be "penalized" if $\boldsymbol{A}_{mean_{i,:}}$ and $\boldsymbol{B}_{mean_{i,:}}$ are far away from each other in the Euclidean space and/or if $\boldsymbol{A}_{mean_{j,:}}$ and $\boldsymbol{B}_{mean_{j,:}}$ are far away from each other in the Euclidean space. $\boldsymbol{A}_{mean_{i,:}}$ indicate the entire row $i$ of $\boldsymbol{A}_{mean}$. To get the distances of genes with identical IDs in $\boldsymbol{A}_{mean}$ and $\boldsymbol{B}_{mean}$, new coordinate sets are defined: $\boldsymbol{C}_{AB_A} = \boldsymbol{C}_{small_{AB}}[:g]$, $\boldsymbol{C}_{AB_B} = \boldsymbol{C}_{small_{AB}}[g:]$, $\boldsymbol{C}_{BA_A} = \boldsymbol{C}_{small_{BA}}[:g]$ and $\boldsymbol{C}_{BA_B} = \boldsymbol{C}_{small_{BA}}[g:]$, where $g$ is the number of genes in $A_{mean}$ and $B_{mean}$. Two vectors, $\boldsymbol{v}_{AB}$ and $\boldsymbol{v}_{BA}$, are calculated by finding the distances between every identically indexed row of $\boldsymbol{C}_{AB_A}$ and $\boldsymbol{C}_{AB_B}$ and every identically indexed row of $\boldsymbol{C}_{BA_A}$ and $\boldsymbol{C}_{BA_B}$, respectively, such that $|\boldsymbol{v}_{AB}| = |\boldsymbol{v}_{BA}| = g$. A vector, $\boldsymbol{v}$, is defined by

$$\boldsymbol{v} = \frac{\boldsymbol{v}_{AB} + \boldsymbol{v}_{BA}}{2}. \quad (6)$$

If the aim is to find clusters of *cluster type 1*, then $v$ is normalized as follows

$$\boldsymbol{v}' = \boldsymbol{v} - min(\boldsymbol{v}), \quad (7)$$

$$\boldsymbol{v}'' = \left| \left( \frac{\boldsymbol{v}'}{(max(\boldsymbol{v}') + 10^{-6})} - 1 \right) \right|. \quad (8)$$

And if clusters of *cluster type 2* is the objective, then $v$ is normalized by

$$\boldsymbol{v}' = \boldsymbol{v} - (min(\boldsymbol{v}) - 10^{-6}), \quad (9)$$

$$\boldsymbol{v}'' = \frac{\boldsymbol{v}'}{max(\boldsymbol{v}')}. \quad (10)$$

In both scenarios, $\boldsymbol{v}$ will be in the range $]0; 1]$, which implies that nothing will be zeroed out by the weighting of $\boldsymbol{v}$. The hyper-similarity matrix is found by weighting row $\boldsymbol{D}_{i,:}$ by element $v_i$ and weighting of column $\boldsymbol{D}_{:,j}$ by element $v_j$ where $i$ is in the range $\{i \in \mathbb{N} \mid 1 \leq i \leq g\}$ and $j$ is in the range $\{j \in \mathbb{N} \mid 1 \leq j \leq g\}$. As a final step, $\boldsymbol{D}$ is normalized by

$$\boldsymbol{D}' = \frac{\boldsymbol{D}}{max(\boldsymbol{D})}, \quad (11)$$

such that all possible gene-gene hyper-similarities are in the range $]0;1]$.

#### Summary - what do hyper-similarity values mean?

When comparing two sets of cells, Scellentor computes a hyper-similarity matrix. The hyper-similarity matrix contains information on how the genes are expressed relative to each other within a set of cells as well as between the compared sets of cells. The main goal of our similarity function is to find genes whose expression patterns are highly similar and conserved within each cell set, but dissimilar between the cell sets (*cluster type 2*). To exemplify this, a high hyper-similarity value (close to 1) between two genes, $gene1$ and $gene2$, implies the following, if e.g. Euclidean distance is used as metric:

i) Both genes express pseudo-timelines in two user-drawn paths, path A and path B; $gene1_A$ and $gene2_A$ are the pseudo-timelines from path A and $gene1_B$ and $gene2_B$ are the pseudo-timelines from path B.

ii) The genes $gene1_A$ and $gene2_A$ are in close proximity to each other and $gene1_B$ and $gene2_B$ are in close proximity to each other.

iii) The distance between $gene1_A$ and $gene2_B$ is large and the distance between $gene1_B$ and $gene2_A$ is large.

iv) The distance between $gene1_A$ and $gene1_B$ is large and the distance between $gene2_A$ and $gene2_B$ is large (weighting by values in vector $\boldsymbol{v}$, Fig. S7).

#### Constrained hierarchical agglomerative clustering

The constrained hierarchical agglomerative clustering iteratively clusters genes together pairwise in order of descending hyper-similarity until all items have been assigned to a cluster or the user-defined threshold has been reached. The user-defined threshold is defined via user-chosen parameters that defines the minimum cluster size and the minimum number of clusters. Per default, the clustering is constrained by the interactions of the PPI network from BioGrid. For the clustering, the constraint means, i) two single genes can only be fused into a cluster if they are neighbors in the network, ii) a single gene needs to neighbor a gene in a cluster before it can be added to the cluster, iii) when merging two clusters, they need to have at least one gene each that are neighbors in the graph. The possible connections of a cluster are equal to the sum of all connections of genes in that cluster.

#### Generating results

When Scellnetor has run a clustering, users can download the main results and all data that were produced in the Scellnetor pipeline. The main results are:

1. Clusters or connected components of genes as PDF files and two edgelists for every cluster in CSV file format where nodes are denoted as both human entrez IDs and human gene symbols
2. One plot per cluster of mean expression of the genes with 95% confidence interval in PDF file format. On the x-axis is "Moving-average-modified number of single cells". The cells are arranged after the variable selected as sorting key. The y-axis shows "Normalized moving-average-modified gene expression". It has been normalized such that the highest value of the concatenation of $\boldsymbol{A}_{mean}$ and $\boldsymbol{B}_{mean}$ is 1 and the smallest is 0. This normalization only serves visualization purposes and is done after the completion of the hyper-similarity calculation.
3. TSV files containing statistically significant GO-terms associated with the clusters. It is generated using GOA-tools[5], which uses Fisher's exact test to calculate p-values and the Benjamini-Hochberg procedure to adjust p-values.

#### Changing the ANNDATA objects and removing noise

After the user has selected a Scanpy-generated plot as template-plot, he/she can choose to remove some of the cells from the ANNDATA object and construct a new one. This functionality has been implemented in case the user only wants to analyze certain groups of cells and perhaps remove noisy cell groups. Here, it makes sense to select a template-plot that is colored by cluster annotations. As mentioned, the clusters you choose will be removed from the data and a new ANNDATA object will be generated that only uses the data that remain. Based on user-defined parameters and user-interaction, pseudotime and novel plots will be generated and stored in the new ANNDATA object.

The new ANNDATA object is now ready for further analysis by the functionalities in the Scellnetor pipeline.

#### Pre-processing of data for Scellentor hematopoiesis study

Using the scRNA-seq data from the 19 clusters defined by ref.[6] (GSE72857), we created an initial count matrix and made sure that we could reproduce their measured cluster-wise average gene expressions before we constructed an ANNDATA object, made cell maps and computed pseudotime. Note, that we could not identify 10 genes from their list with cluster-wise average gene expressions. These anonymous genes were omitted from our analysis. This gave us a total of 2730 single cells that expressed 3451 genes. We pre-processed the count matrix the same way as in the study by ref.[7], with the exception of using the top 1726 ($3451/2 \approx 1726$) most variable genes instead of the top 1000.

#### Pre-processing of data for Scellentor exhausted CD8 T-cell study

Using the scRNA-seq data from the five clusters they identified in ref.[8] (GSE137007) we generated an ANNDATA object. The data was already pre-processed using the Seurat package. Additionally, we computed a diffusion map and calculated pseudo time. The “start cell” for the pseudotime inference was the cell in the progenitor cluster (cluster 1, Fig. 3a) that was furthest away from all progenies.
