## Supplementary figure legends for "Comparative single-cell trajectory network enrichment identifies pseudo-temporal systems biology patterns in hematopoiesis and CD8 T-cell development"

**Figure S1. Cell map based on force-directed graph drawing of cell map.** Cell map shows localization of single cells from the study ref. [1]. On the right is shown the color codes for the cluster annotations.

**Figure S2. Clusters and mean pseudo-timelines and 95 % confidence intervals from Scellentor hematopoiesis study. a-d** Gene modules and plots are based on cells from ref. [1]. Mean pseudo-timelines and 95 % confidence intervals of the genes in the Scellnetor clusters 4-7. The q-values below every plot are based on Wilcoxon signed-rank tests where the average expression over time of the clustered genes from the two distinct differentiation paths is compared. **e-h** Connected subnetworks of genes from clusters 4-7. The q-value is shown in the grey boxes near every cluster are based on a p-value from a Mann-Whitney *U* test, where all possible hyper-similarity values are compared with the values of the clusters. All p-values shown on the figure are corrected via the Benjamini-Hochberg Procedure to produce q-values.

**Figure S3. Average expression of genes in Scellnetor clusters. a-g** Red line indicate clusters 15-17 and green line indicate clusters 1-6 from ref. [1]. **a-g** Correspond to Scellnetor hematopoiesis clusters 1-7. Along the first axes are names of genes in the Scellnetor hematopoiesis clusters. Along the second axes are shown the average gene expressions. P-values and q-values are shown below every plot. P-values are from Wilcoxon signed-rank tests where values from red lines are compared to values from green lines. Each p- and q-value pair are based on the values from the plots above them. Q-values are found using the Benjamini-Hochberg Procedure.

**Figure S4. Average expression of genes in Scellnetor clusters - cells from paths. a-g** Red line indicate clusters 15-17 and green line indicate clusters 1-6 from ref. [1]. **a-g** Correspond to Scellnetor hematopoiesis clusters 1-7. Along the first axes are names of genes in the Scellnetor hematopoiesis clusters. Along the second axes are shown the average pre-processed gene expressions. P-values and q-values are shown below every plot. P-values are from Wilcoxon signed-rank tests where values from red lines are compared to values from green lines. Each p- and q-value pair are based on the values from the plots above them. Q-values are found using the Benjamini-Hochberg Procedure.

**Figure S5. Stem cell area on cell map based on force-directed graph drawing.** Yellow cells constitute the stem cell area comprised of cluster 7-10 from ref. [1]. Purple cells are all other cells. Blue point is the average position of the stem cell area (shown as a square). Orange point is the cell closest to the average position of the stem cell area and the "start cell" of the pseudotime computation (shown as a square). On the right is shown a box with color coding of the cells.

**Figure S6. Workflow of the Scellnetor pipeline.** **1** User uploads a H5AD file containing scRNA-seq data or tries a test set. **2** User chooses desired Template-plot. **3** Selection of cells on Canvas-plot that represents differentiation trajectories. User can choose between drawing paths (**3a**) or selecting clusters (**3b**). **4** User defines cluster parameters such as linkage type, distance metric, number of clusters and minimum sizes of clusters. **5** Scellentor extracts selected cells and dimensionality reduce gene expression patterns. This is followed by a calculation of a hyper-similarity matrix (if comparing two sets of cells). Afterwards, Scellnetor performs a constrained hierarchical clustering based on modified expression data from the selected cells. 6 User can inspect and download the results from the Scellnetor clustering.

**Figure S7. Computation of hyper-similarity matrix - extended figure.** **1** Two paths are drawn on the canvas-plot, red path and green path. Messenger-RNA expression data are extracted from selected cells and used to create expression matrices – one for each drawn path. The expression matrices are dimensionality reduced with UMAP, such that every pseudo-timeline of genes in the two sets are converted to 2D coordinates in the Euclidean space. **2** The 2D coordinates from the red set and the green set are concatenated along the first axis. The concatenated coordinate sets are used to produce a distance matrix, $\boldsymbol{D}$, which is normalized by division of its max value. The distances between genes with identical IDs, but from different sets, are stored in the vector, $\boldsymbol{v}$ .The diagonal of the upper right quadrant of $\boldsymbol{D}$ is zeroed-out (top, right). Triangular matrices of the distance matrix' quadrant I, II and IV (as in cartesian coordinate system) are extracted. Matrix $\boldsymbol{D}_{rr}$ corresponds to a distance matrix of "red set vs. red set", matrix $\boldsymbol{D}_{gg}$ corresponds to a distance matrix of "green set vs. green set", matrix $\boldsymbol{D}_{rg}$ corresponds to a distance matrix of "red set vs. green set" and matrix $\boldsymbol{D}_{gr}$ corresponds to a distance matrix of "green set vs. red set". **3** Matrices $\boldsymbol{D}_{rr}$ and $\boldsymbol{D}_{gg}$ are modified such that genes with the shortest distances have the highest values and vice versa. The matrix $\boldsymbol{D}_{gr}$ is transposed and the Hadamard product of all triangular matrices is found (bottom, left). This produces the hyper-similarity matrix, $\boldsymbol{D}_{hyp}$. The rows and columns of $\boldsymbol{D}_{hyp}$ are weighted by the values of $\boldsymbol{v}$ (bottom, right).
