## Supplementary figures for "Comparative single-cell trajectory network enrichment identifies pseudo-temporal systems biology patterns in hematopoiesis and CD8 T-cell development"

### Figure S1

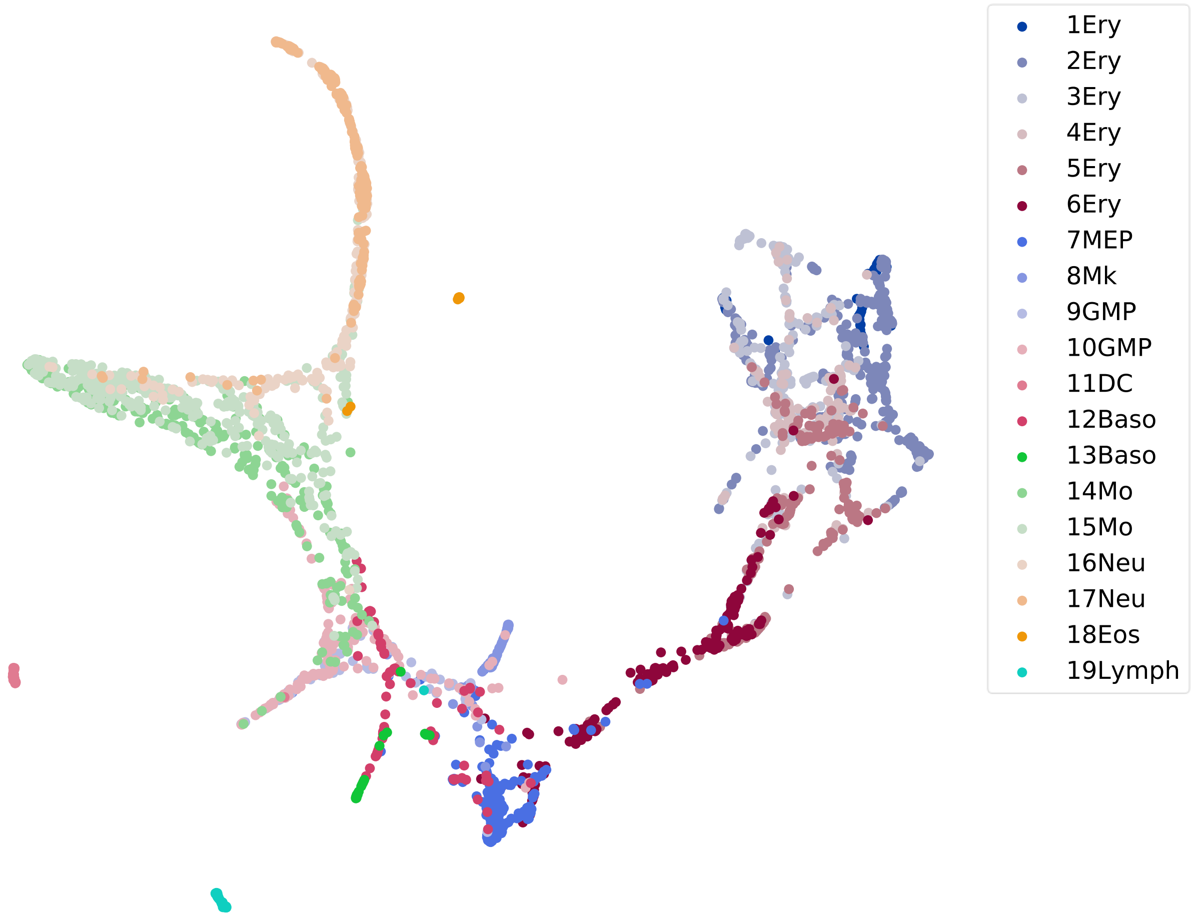
1Ery
2Ery
3Ery
4Ery
5Ery
6Ery
7MEP
8Mk
9GMP
10GMP
11DC
12Baso
13Baso
14Mo
15Mo
16Neu
17Neu
18Eos
19Lymph

### Figure S2

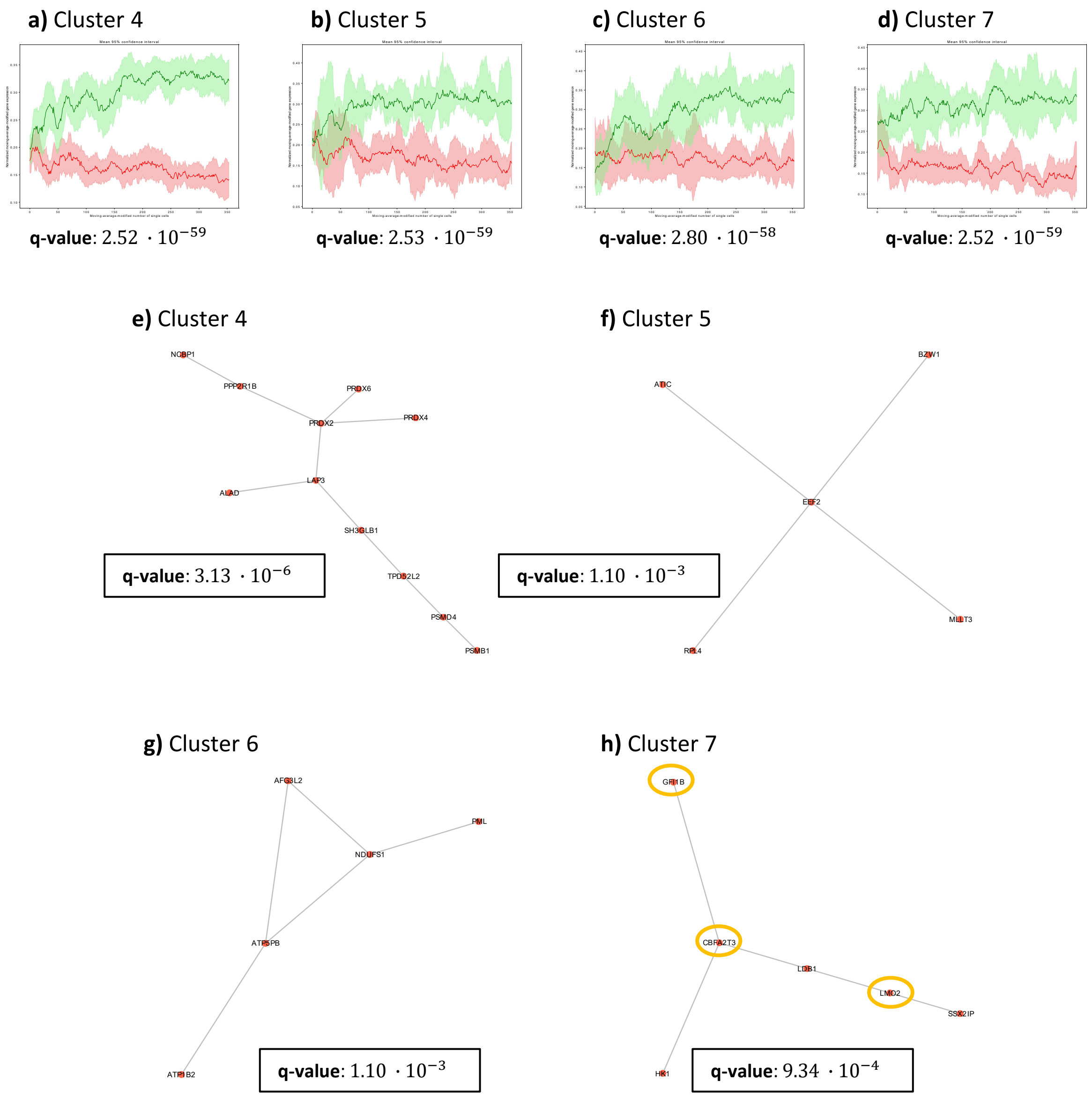

a) Cluster 4
q-value: $2.52 \cdot 10^{-59}$
b) Cluster 5
q-value: $2.53 \cdot 10^{-59}$
c) Cluster 6
q-value: $2.80 \cdot 10^{-58}$
d) Cluster 7
q-value: $2.52 \cdot 10^{-59}$
e) Cluster 4
NCBP1
PPP2R1B
PRDX6
PRDX2
PRDX4
LAP3
ALAD
SH3GLB1
TPD52L2
PSMD4
PSMB1
q-value: $3.13 \cdot 10^{-6}$
f) Cluster 5
BZW1
ATIC
EEF2
MLLT3
RPL4
q-value: $1.10 \cdot 10^{-3}$
g) Cluster 6
AFG3L2
PML
NDUFS1
ATP5PB
ATP1B2
q-value: $1.10 \cdot 10^{-3}$
h) Cluster 7
GFI1B
CBFA2T3
LDB1
LMO2
SSX2IP
HK1
q-value: $9.34 \cdot 10^{-4}$

### Figure S3

**a)** Cluster 1 **b)** Cluster 2 **c)** Cluster 4 **d)** Cluster 5 **e)** Cluster 6 **f)** Cluster 7

Average gene expression of compared clusters according to Paul et al. (2015)

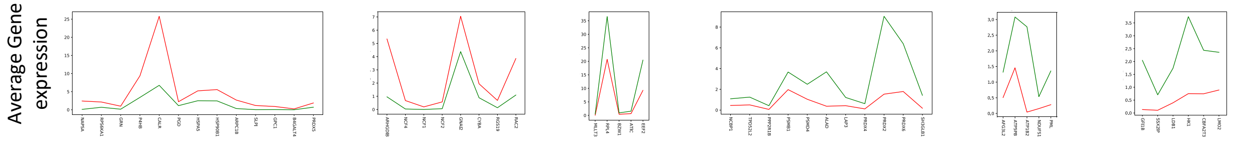


Genes

| | a) Cluster 1 | b) Cluster 2 | c) Cluster 4 | d) Cluster 5 | e) Cluster 6 | f) Cluster 7 |
|---|---|---|---|---|---|---|
| P-value | $1.47 \cdot 10^{-3}$ | 0.012 | $3.35 \cdot 10^{-3}$ | 0.043 | 0.043 | 0.028 |
| Q-value | $5.16 \cdot 10^{-3}$ | 0.021 | $7.81 \cdot 10^{-3}$ | 0.043 | 0.043 | 0.039 |

**g)** Cluster 3

Average gene expression of compared clusters according to Paul et al. (2015)

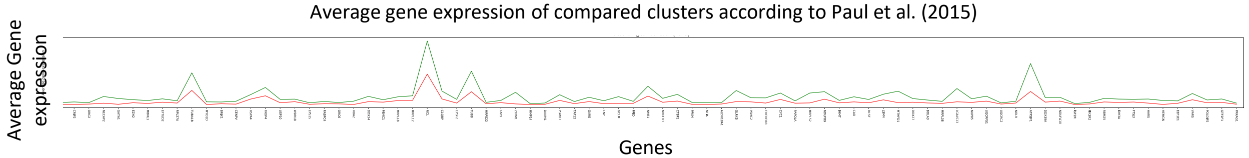


Genes

P-value $5.36 \cdot 10^{-15}$

Q-value $3.75 \cdot 10^{-14}$

### Figure S4

**a)** Cluster 1 **b)** Cluster 2 **c)** Cluster 4 **d)** Cluster 5 **e)** Cluster 6 **f)** Cluster 7

Average gene expression of compared clusters according to Paul et al. (2015)

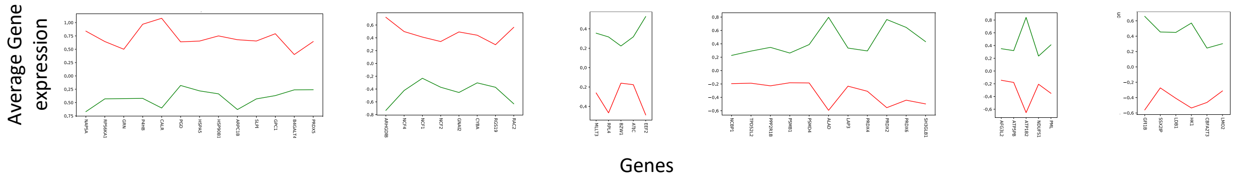


| | a) Cluster 1 | b) Cluster 2 | c) Cluster 4 | d) Cluster 5 | e) Cluster 6 | f) Cluster 7 |
|---|---|---|---|---|---|---|
| P-value | $1.47 \cdot 10^{-3}$ | 0.012 | $3.35 \cdot 10^{-3}$ | 0.043 | 0.043 | 0.028 |
| Q-value | $5.16 \cdot 10^{-3}$ | 0.021 | $7.81 \cdot 10^{-3}$ | 0.043 | 0.043 | 0.039 |

**g)** Cluster 3

Average gene expression of compared clusters according to Paul et al. (2015)

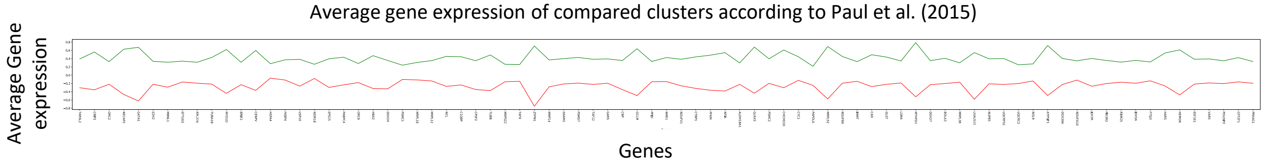


| | g) Cluster 3 |
|---|---|
| P-value | $5.36 \cdot 10^{-15}$ |
| Q-value | $3.75 \cdot 10^{-14}$ |

### Figure S5

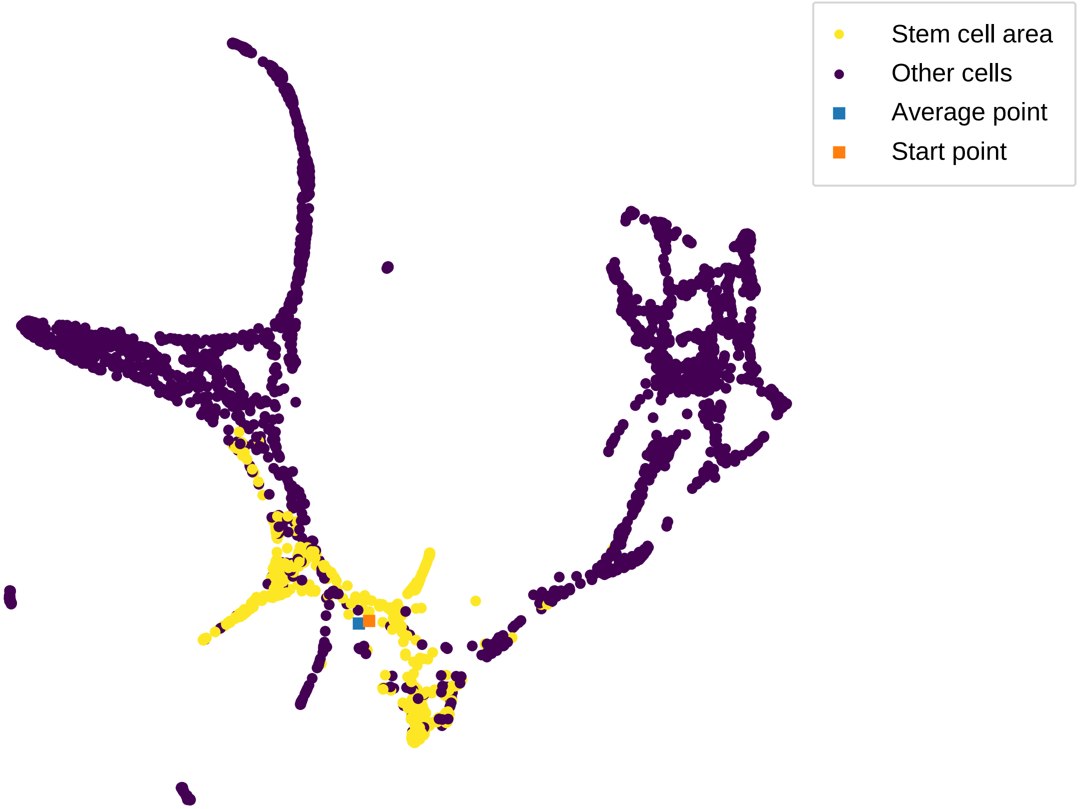

Figure S6

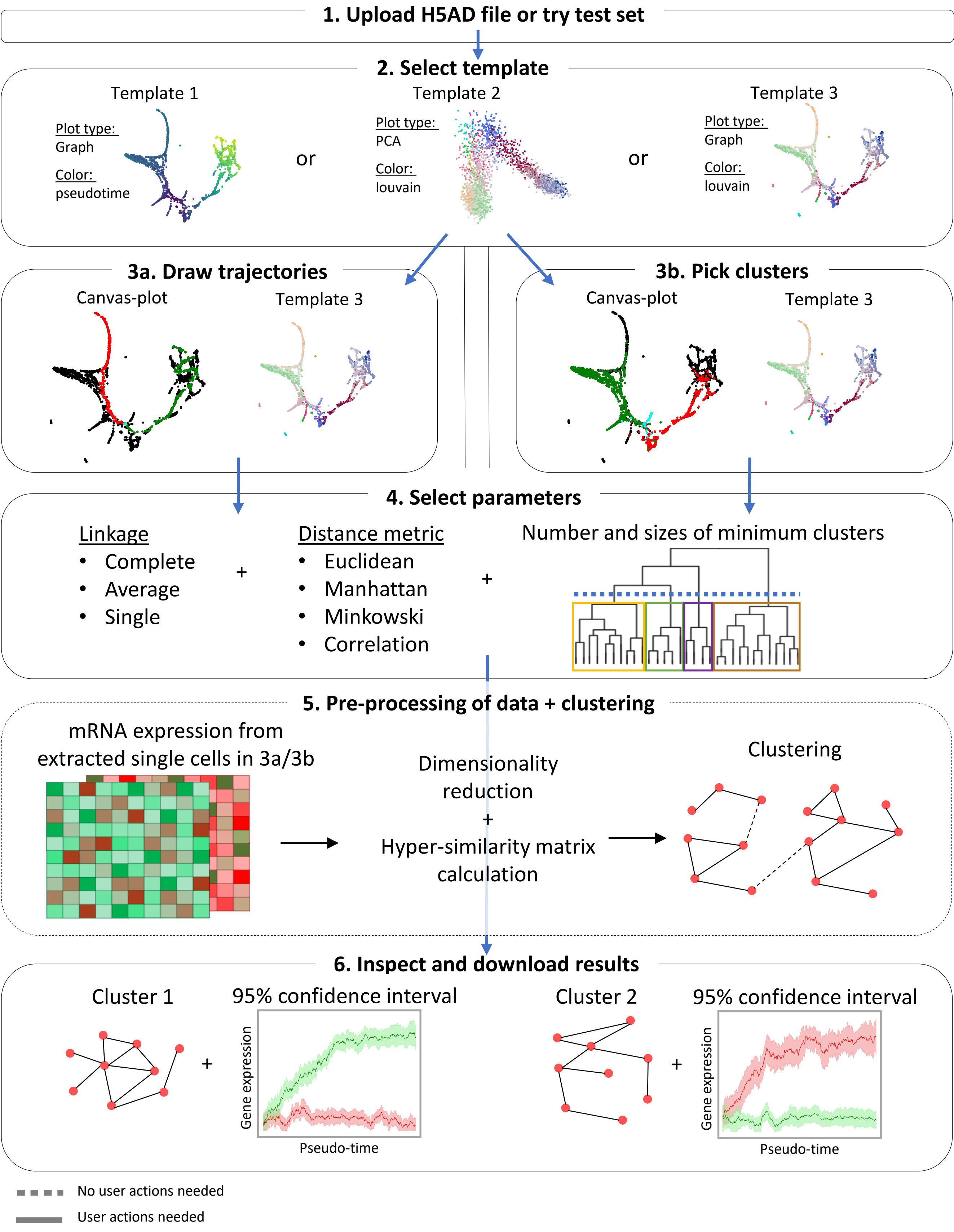

1. Upload H5AD file or try test set
2. Select template
Template 1
Plot type:
Graph
Color:
pseudotime
or
Template 2
Plot type:
PCA
Color:
louvain
or
Template 3
Plot type:
Graph
Color:
louvain
3a. Draw trajectories
Canvas-plot
Template 3
3b. Pick clusters
Canvas-plot
Template 3
4. Select parameters
Linkage
Complete
Average
Single
+
Distance metric
Euclidean
Manhattan
Minkowski
Correlation
+
Number and sizes of minimum clusters
5. Pre-processing of data + clustering
mRNA expression from extracted single cells in 3a/3b
Dimensionality reduction
+
Hyper-similarity matrix calculation
Clustering
6. Inspect and download results
Cluster 1
+
95% confidence interval
Gene expression
Pseudo-time
Cluster 2
+
95% confidence interval
Gene expression
Pseudo-time
No user actions needed
User actions needed

Figure S7

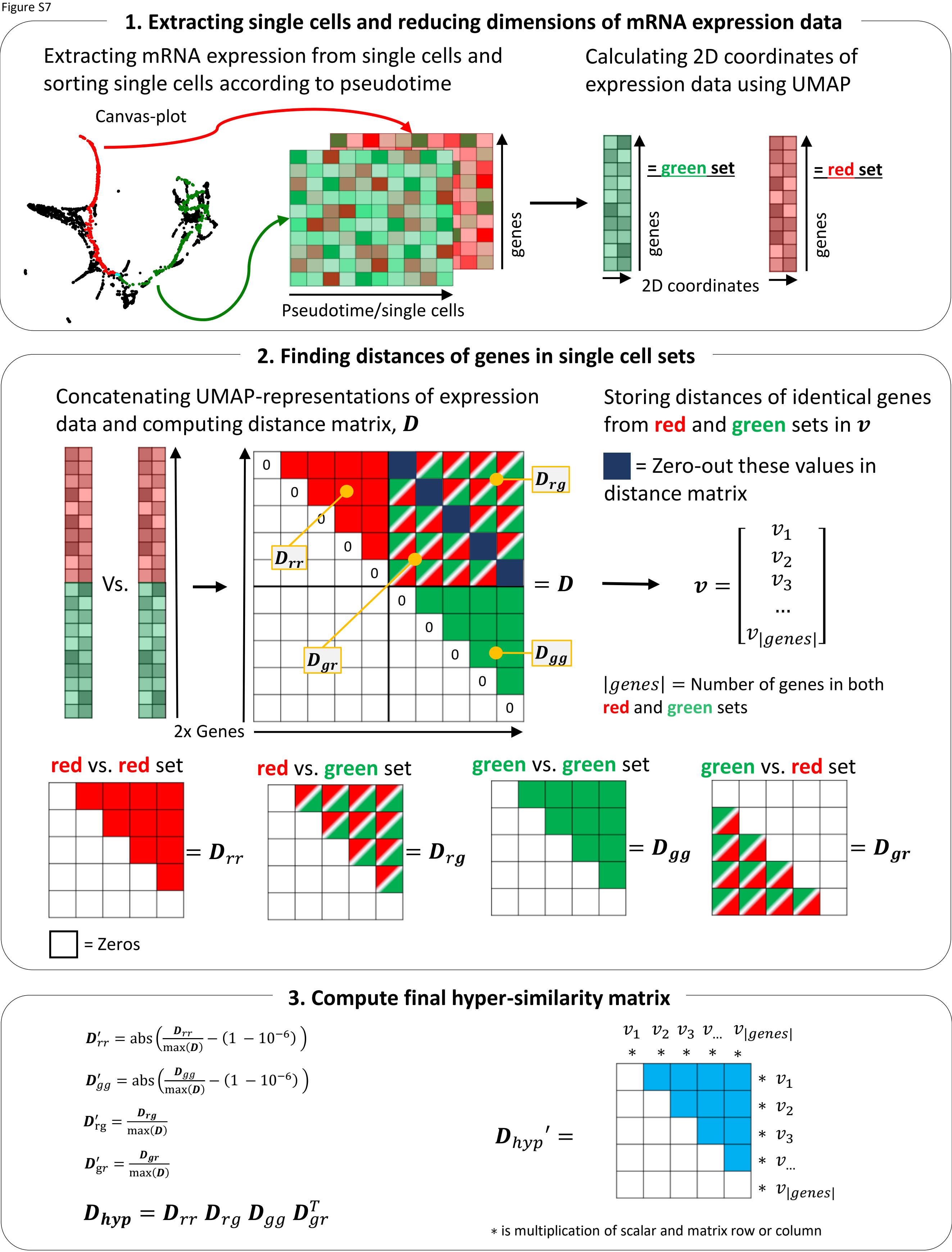

1. Extracting single cells and reducing dimensions of mRNA expression data
Extracting mRNA expression from single cells and sorting single cells according to pseudotime
Canvas-plot
genes
Pseudotime/single cells
Calculating 2D coordinates of expression data using UMAP
= green set
genes
= red set
genes
2D coordinates
2. Finding distances of genes in single cell sets
Concatenating UMAP-representations of expression data and computing distance matrix, $\boldsymbol{D}$
Vs.
2x Genes
0
0
0
0
0
0
0
0
0
0
$D_{rg}$
$D_{rr}$
$D_{gr}$
$D_{gg}$
$= \boldsymbol{D}$
Storing distances of identical genes from red and green sets in $\boldsymbol{v}$
= Zero-out these values in distance matrix
$\boldsymbol{v} = [v_1, v_2, v_3, \dots, v_{|genes|}]$
$|genes|$ = Number of genes in both red and green sets
red vs. red set
$= D_{rr}$
red vs. green set
$= D_{rg}$
green vs. green set
$= D_{gg}$
green vs. red set
$= D_{gr}$
= Zeros
3. Compute final hyper-similarity matrix
$D'_{rr} = \text{abs}\left(\frac{D_{rr}}{\max(\boldsymbol{D})} - (1 - 10^{-6})\right)$
$D'_{gg} = \text{abs}\left(\frac{D_{gg}}{\max(\boldsymbol{D})} - (1 - 10^{-6})\right)$
$D'_{rg} = \frac{D_{rg}}{\max(\boldsymbol{D})}$
$D'_{gr} = \frac{D_{gr}}{\max(\boldsymbol{D})}$
$D_{hyp} = D_{rr}\ D_{rg}\ D_{gg}\ D_{gr}^{T}$
$v_1$ $v_2$ $v_3$ $v_{\dots}$ $v_{|genes|}$
* * * * *
$D_{hyp}' =$
* $v_1$
* $v_2$
* $v_3$
* $v_{\dots}$
* $v_{|genes|}$
* is multiplication of scalar and matrix row or column
